# CORTADO: Hill Climbing Optimization for Cell-Type Specific Marker Gene Discovery

**DOI:** 10.1101/2024.12.23.630040

**Authors:** Musaddiq K Lodi, Leiliani Clark, Satyaki Roy, Preetam Ghosh

**Affiliations:** Integrative Life Sciences, Virginia Commonwealth University, Richmond, VA, United States of America; Center for Biological Data Science, Virginia Commonwealth University, Richmond, VA, United States of America; Department of Mathematical Sciences, University of Alabama in Huntsville, Huntsville, AL, United States of America; Department of Computer Science, Virginia Commonwealth University, Richmond, VA, United States of America

**Keywords:** Single-cell RNA-seq, marker gene discovery, hill climbing, optimization, cellular heterogeneity

## Abstract

The advent of single-cell RNA sequencing (scRNA-seq) has greatly enhanced our ability to explore cellular heterogeneity with high resolution. Identifying subpopulations of cells and their associated molecular markers is crucial in understanding their distinct roles in tissues. To address the challenges in marker gene selection, we introduce CORTADO, a computational framework based on hill-climbing optimization for the efficient discovery of cell-type-specific markers. CORTADO optimizes three critical properties: differential expression in the clusters of interest, distinctiveness in gene expression profiles to minimize redundancy, and sparseness to ensure a concise and biologically meaningful marker set. Unlike traditional methods that rely on ranking genes by p-values, CORTADO incorporates both differential expression metrics and penalties for overlapping expression profiles, ensuring that each selected marker uniquely represents its cluster while maintaining biological relevance. Its flexibility supports both constrained and unconstrained marker selection, allowing users to specify the number of markers to identify, making it adaptable to diverse analytical needs and scalable to datasets with varying complexities. To validate its performance, we apply CORTADO to several datasets, including the DLPFC 151507 dataset, the Zeisel mouse brain dataset, and a peripheral blood mononuclear cell dataset. Through enrichment analysis and examination of spatial localization-based expression, we demonstrate the robustness of CORTADO in identifying biologically relevant and non-redundant markers in complex datasets. CORTADO provides an efficient and scalable solution for cell-type marker discovery, offering improved sensitivity and specificity compared to existing methods.

## Introduction

The emergence of single-cell RNA sequencing (scRNA-seq) has enabled researchers to explore transcriptional processes at single-cell resolution. This technology has been instrumental in identifying rare cell types, evaluating cellular heterogeneity, and measuring variations between individual cells [1]. Single-cell biology focuses on characterizing cellular heterogeneity by uncovering distinct sub-populations [2]. Many methods have been developed to perform this task, ranging from unsupervised clustering, to reference dataset-based cell annotation [3–5]. Identifying marker genes is a key step for subtyping cell populations, regardless of the method used for cell population identification.

Marker genes, while variably defined in the literature without a universal consensus, serve the essential purpose of distinguishing between different cell populations. A recent benchmarking work of marker gene selection methods notably defined marker genes as a subset of the differentially expressed genes of a cluster that have the most relevance to the cell type’s biological function [6]. The most common tool for marker gene identification is log fold change-based statistical testing, such as the Wilcoxon Rank Sum and t-Test [7] and have been incorporated in widely used scRNA-seq workflows [7, 8].

Calculating the differential expression of a gene alone may not reliably identify an accurate set of marker genes for a specific cluster. For instance, a gene might show high expression in the target cell type but also moderately high expression in another cell type, with low expression in the rest. Existing approaches leveraging expression-based statistical methods might still identify this gene as a marker, even though it is not exclusive to the target cell type, highlighting the need for a more thorough assessment of gene expression patterns to ensure proper marker gene selection. Several marker gene selection methods have emerged in recent times. One method, RankCorr, identifies marker genes by calculating the Spearman correlation with the cluster indicator vector for each fixed cluster and using that information to identify appropriate markers [9]. Another technique, scMAGS, utilizes differential expression or variability to calculate cluster-specific marker scores and cluster validity indices, Silhouette index or Calinski-Harabasz index to select such marker genes [10]. COSG introduced the importance of considering the cosine similarity profile of certain genes in marker selection [11].

Current approaches to marker gene selection often fail to provide a holistic view of differential gene expression while adequately addressing overlap or redundancy in gene expression profiles [12–14]. Additionally, existing methods typically rely on ranking genes by p-adjusted values, failing to identify a concise and biologically meaningful set of marker genes. To address these challenges, we present CORTADO (hill Climbing OptimizaTion foR cell-type specific mArker gene DiscOvery), a framework that selects markers by optimizing three critical properties: differential expression in the clusters of interest, distinctiveness in their expression profiles (or non-redundancy) concerning other markers, and sparseness, i.e., minimizing the number of selected genes (as illustrated in Figure 1). By incorporating differential expression metrics and penalizing gene pairs with overlapping profiles, CORTADO ensures that each selected marker uniquely characterizes its cluster of interest while remaining biologically interpretable.

**Fig 1.**
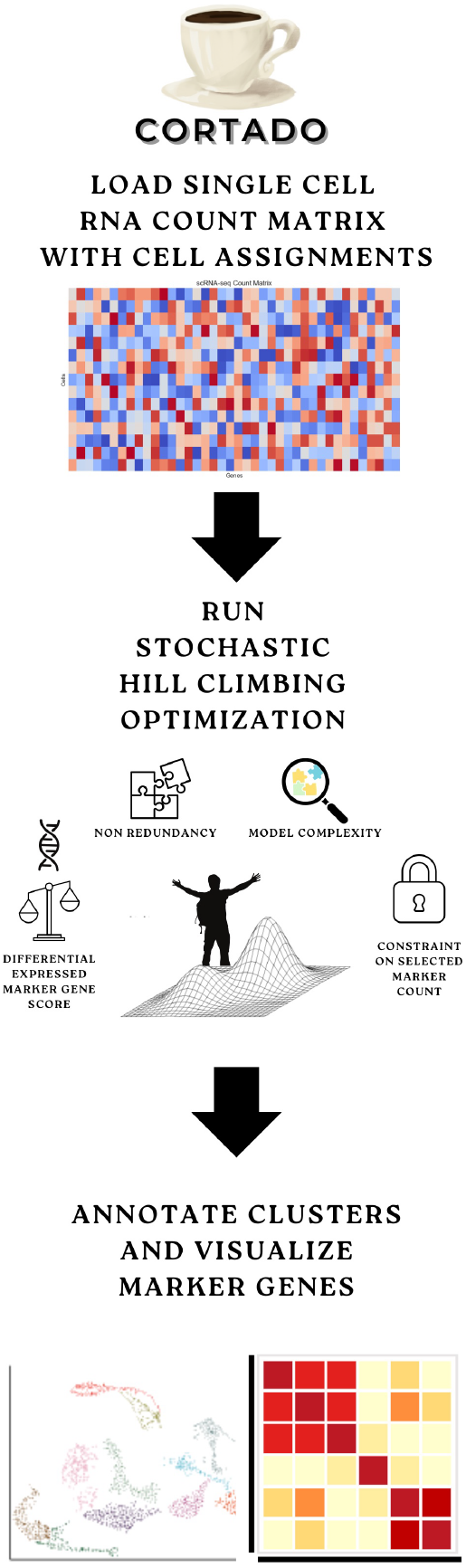
Overview of CORTADO workflow. CORTADO is a marker gene selection framework with three main steps. First, a single cell genomics count matrix is loaded as a Scanpy object and undergoes standard preprocessing. Then, the stochastic hill optimization is run, with four key components taken into consideration when selecting genes: Differential expressed gene score, non-redundancy based on cosine similarity, a penalization of selecting too many genes, and a constraint parameter to select a user-defined amount of genes. Then, the genes can be visualized through expression heatmaps, and contextualized using relevant literature and gene set enrichment analyses.

CORTADO also introduces flexibility that many existing methods lack, supporting both constrained and unconstrained scenarios for marker selection. In the constrained scenario, users can specify the number of marker genes to be identified, enabling tighter control over the size of the gene set. This flexibility makes CORTADO adaptable to diverse analytical needs and scalable to datasets with varying complexity. Furthermore, the hill-climbing optimization framework underpinning CORTADO efficiently balances the need for non-redundant and sparse marker sets, addressing redundancy in gene expression profiles without compromising the biological relevance of the selected markers.

Through enrichment analyses and the examination of spatial localization-based expression, we establish the biological relevance and uniqueness of CORTADO-selected markers through three case studies. On the DLPFC 151507 dataset, derived from spatial transcriptomics data of the dorsolateral prefrontal cortex, CORTADO identifies GFAP, a well-known significant marker of astrocytes. In the Zeisel mouse brain dataset, containing 3005 cells from hippocampal tissue, CORTADO highlights unique genes associated with the “Modulation of the Chemical Synaptic Transmission” pathway, underscoring its capacity for non-redundant selection in neuronal communication pathways. Analysis of a peripheral blood mononuclear cell dataset reveals that the selected markers are enriched in pathways such as viral transcription and immune response, showcasing CORTADO’s ability to identify significant genes with clinical relevance. CORTADO is implemented in Python and available for installation at https://www.github.com/lodimk2/cortado.

## Results

### CORTADO Overview

CORTADO, implemented using a *stochastic hill optimization* framework, was evaluated for its ability to select cluster-specific marker genes across diverse scRNA-seq datasets. Starting with a raw count matrix and a metadata file containing clustering assignments, the data was processed into an AnnData object [7]. Marker genes and their cosine similarity scores were calculated for each user-specified cluster of interest. Additionally, CORTADO includes functionality for identifying markers across all clusters within a dataset. As depicted in Figure 1, it operates under an objective function comprising three weighted terms (Refer to *Methods* for more details). The first term, *c*_1_, prioritizes genes with strong differential expression in the cluster of interest compared to other clusters. The second term, *c*_2_, penalizes the selection of gene pairs with high cosine similarity, thereby promoting marker diversity within the cluster of interest. The third term, *c*_3_, discourages the selection of an excessive number of genes, enforcing sparsity in the final marker set. Each term is weighted by a user-defined parameter *λ*_*i*_ (*i* = 1, 2, 3), with larger *λ* values assigning greater importance to the corresponding term. Overall, the objective function is: *λ*_1_*c*_1_ −*λ*_2_*c*_2_ −*λ*_3_*c*_3_.

Based on a sensitivity analysis using simulated datasets with well-separated and overlapping clusters, we recommend parameter values of *λ*_1_ = 0.9 and *λ*_2_ = 0.1 for most scRNA-seq datasets. Details of this analysis are provided in the Supplementary Materials. To determine the effect of *λ*_3_ on the number of selected genes by CORTADO when the model is unconstrained, we show that increasing the value of *λ*_3_ decreases the number of genes selected, across clusters in the Zeisel Mouse Brain Dataset [15]. To benchmark the performance of CORTADO, we tested its ability to identify marker genes across multiple scRNA-seq datasets, including the Zeisel Mouse Brain dataset, the Baron Pancreas dataset, the Muraro Pancreas dataset, and the PBMC3K dataset [15–18]. Furthermore, we evaluated its utility in a disease context using a Basal Cell Carcinoma (BCC) dataset [19].

### CORTADO selects more differentiated markers based on gene expression and gene similarity

To benchmark CORTADO, we devised two scores to assess marker gene strength: the *log ratio difference* and the *median cosine similarity*. The log ratio difference measures the difference in expression of selected marker genes in the cluster of interest versus all other clusters, while cosine similarity evaluates the similarity of genes within the cluster of interest relative to other clusters. (Refer to *Methods* for more details)

Pullin and McCarthy [6] proposed log ratio difference as a key indicator, assuming strong markers have high expression differences in the target cluster. However, simple differential expression may not fully capture marker gene quality, as markers should distinguish cell subpopulations beyond expression alone [6]. Cosine similarity, a scale-free measure, complements this by identifying genes with diverse profiles outside the target cluster [11]. Importantly, the absence of cosine similarity between genes enables the selection of markers with minimal overlap in expression profiles, ensuring better distinction across subpopulations. Thus, we argue that robust marker selection requires both metrics.

CORTADO was compared to four baseline methods across seven datasets using these metrics. To rank the performance of each method, we took the median of the performance metrics. For the log ratio metric, a higher median log ratio corresponded to a better rank. For the cosine similarity metric, a median lower cosine similarity corresponded a better rank (a rank of 1 is considered the best rank). Figure 2 illustrates these comparisons, where the points represent the average rank of each method across evaluations, while the error bars indicate the standard deviation of median scores. In Figure 2A, CORTADO achieves the top rank with minimal variability, outperforming other methods. Figure 2B shows COSG leading in cosine similarity difference, with CORTADO consistently ranking second. COSG and CORTADO exhibited no rank variability in this metric. These results highlight CORTADO’s strengths. It excels in selecting genes with high log-ratio differences, ensuring strong expression in the cluster of interest and low expression elsewhere. Additionally, its strong performance in cosine similarity difference demonstrates its ability to identify diverse markers while minimizing overlap in expression profiles.

**Fig 2.**
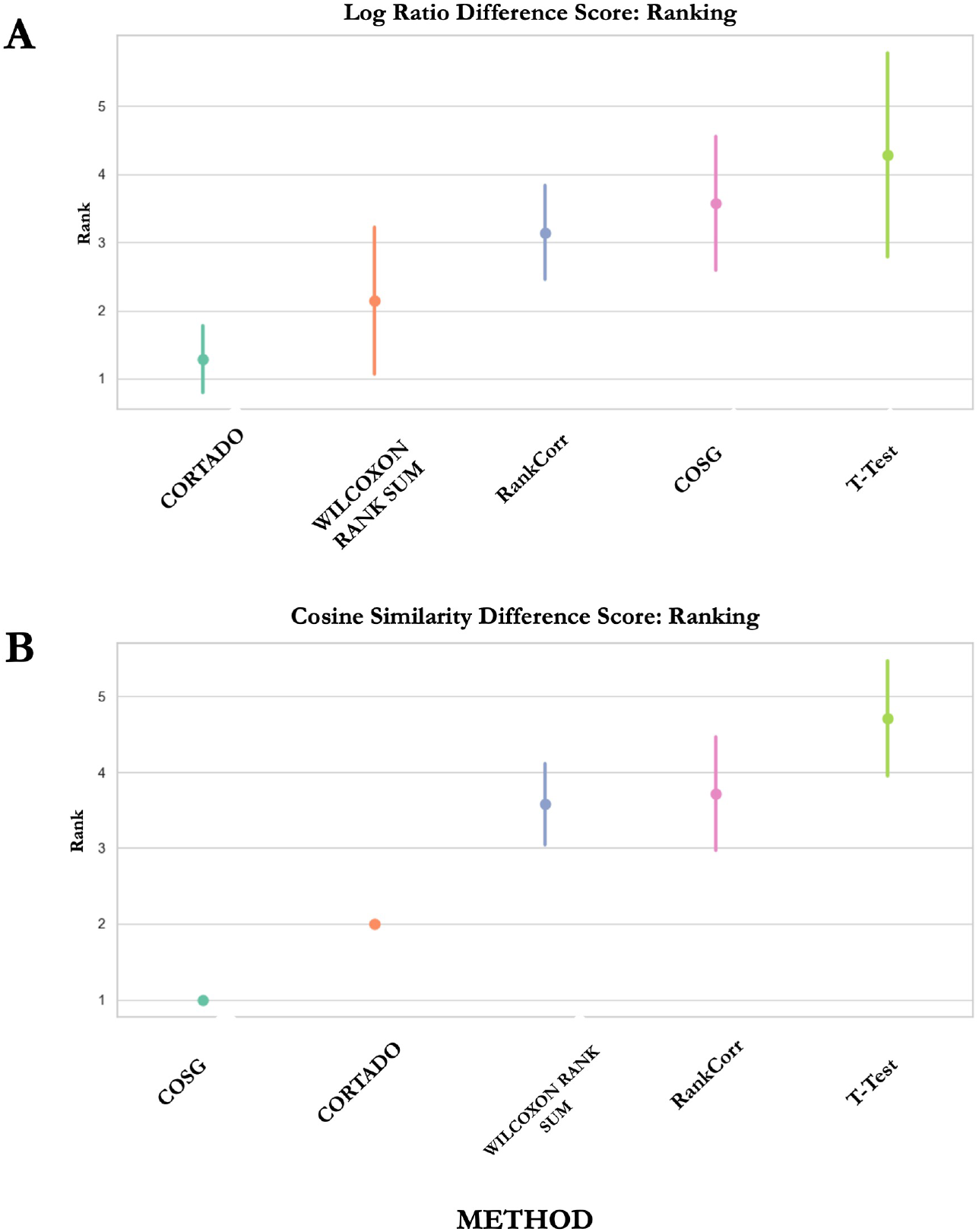
Performance ranking across 7 datasets based on log-ratio difference score and cosine similarity score. A lower number corresponds to a better rank. A) Ranking of baseline methods across datasets based on log-ratio difference score. B) Ranking of baseline methods across datasets based on cosine similarity difference score.

### CORTADO can distinguish high fidelity marker genes in the Zeisel Mouse Brain Dataset compared to baseline methods

The Zeisel mouse brain dataset contains 3005 cells from mouse hippocampus tissue, with 7 distinct clusters [15]. This dataset is commonly used for scRNAseq benchmarking studies, as the clusters are biologically relevant and well separated [9], making it relevant for evaluating marker selection methods. The heatmap in Figure 3B depicts the expression of the top 5 selected marker genes for each cluster across all methods. This result is a very clear indicator of genes that may have high expression in more than one cell type. While it is improbable to expect that selected markers will only have expression in the cluster of interest, strong markers should generally have high expression in its intended cluster, and low expression in all other clusters. While COSG and RankCorr select the most distinct genes based on the marker gene score, CORTADO follows a similar pattern by selecting well distinguished genes, particularly in the astrocyte-ependymal, endothelial-mural, and microglia clusters. From this analysis, we see in 3A that the markers selected by CORTADO, RankCorr, and COSG follow a pattern where the markers selected have high expression in one cluster, and negligible expression in other clusters. The Scanpy implementations of the Wilcoxon Rank Sum test and t-test appear to have selected markers that have consistent expression in multiple clusters [7]. COSG in particular selected genes that are distinguished in their cluster by expression value.

**Fig 3.**
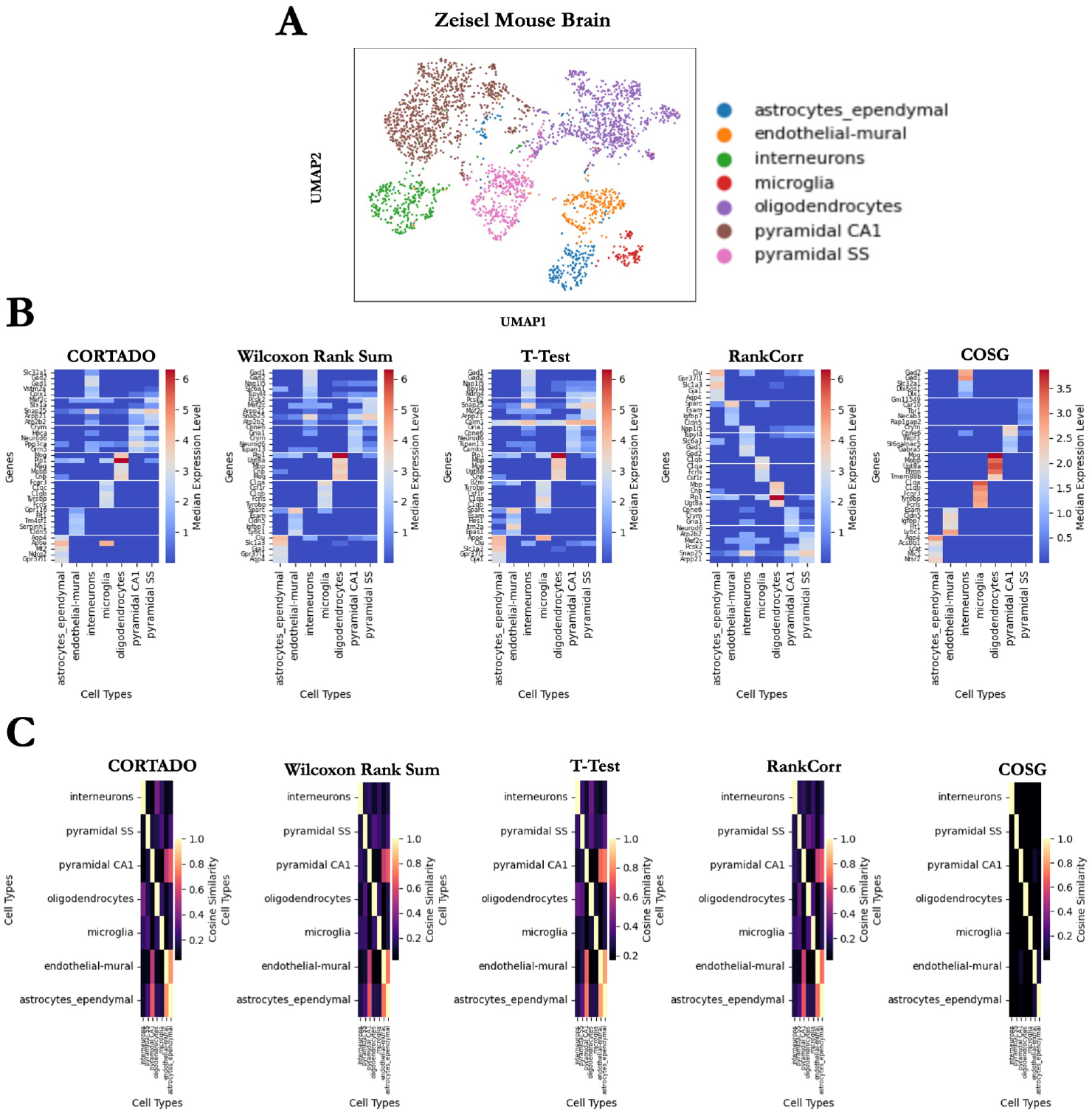
CORTADO analysis on the Zeisel Mouse Brain Dataset. A) Median expression level for top 5 selected markers per method across cell types. B) Cosine similarities between cell-type median gene expression for the top 5 selected markers per method. C) UMAP Visualization for cell types in the Zeisel Mouse Brain dataset.

The heatmap in Figure 3C shows the cosine similarity between markers selected in each method across clusters. The cosine similarity for the heatmap was calculated by averaging the expression across markers for each cluster and then computing the cosine similarity of the average expression vector. Strong markers selected by cosine similarity separation have low cosine similarity across clusters. CORTADO and COSG select markers with low cosine similarity across clusters, while other methods choose markers with high cosine similarity. The results in Figure 3B and Figure 3C confirm these findings from the quantitative benchmarking in a practical use case.

In the following experiment, we elucidate CORTADO’s capability to identify unique and biologically relevant genes that are often missed by other approaches. A gene is deemed *unique* if it is selected by only a single approach for a given cluster. Figure 4 shows that CORTADO and COSG select the highest number of unique genes aggregated across clusters. To verify their biological relevance, we used the Pyramidal CA1 cell cluster. Pyramidal CA1 cells are a type of neuron that processes sensory and motor cues [20]. Pyramidal CA1 neurons are also the most abundant neurons in the brain [21]. The unique genes selected by CORTADO to be markers for the Pyramidal CA1 cluster were Grm5, Hpca, and Ppp3ca. Furthermore, we ran a GO Biological Process (BP) enrichment using the EnrichR online tool [22] [23]. We down-selected significant pathways based on FDR Corrected p-value satisfying a 0.05 cutoff threshold, and sorted by p-value after applying the cutoff. The results of this analysis are depicted in the bar plot in Table 1. We see from this figure that the pathways being selected are related to signaling overall. Of particular interest is the Modulation of the Chemical Synaptic Transmission pathway; chemical synaptic transmission is imperative to neuronal communication, as this pathway facilitates the release of neurotransmitters in the synapse [24].

**Table 1.** Enrichment Analysis Results with GO Terms and Descriptions for Selected Markers Zeisel Mouse Brain Dataset: Pyramidal CA1 Cells.

| GO Term | Description |
| --- | --- |
| GO:0019722 | Calcium-Mediated Signaling |
| GO:0050804 | Modulation of Chemical Synaptic Transmission |
| GO:0010359 | Regulation of Anion Channel Activity |
| GO:0060993 | Kidney Morphogenesis |
| GO:0043403 | Skeletal Muscle Tissue Regeneration |
| GO:0046877 | Regulation of Saliva Secretion |
| GO:0090184 | Positive Regulation of Kidney Development |
| GO:1905665 | Positive Regulation of Calcium Ion Import Across Plasma Membrane |
| GO:0048741 | Skeletal Muscle Fiber Development |
| GO:0070262 | Peptidyl-Serine Dephosphorylation |
| GO:0033173 | Calcineurin-NFAT Signaling Cascade |
| GO:0097205 | Renal Filtration |
| GO:1905664 | Regulation of Calcium Ion Import Across Plasma Membrane |
| GO:0051047 | Positive Regulation of Secretion |
| GO:0097720 | Calcineurin-Mediated Signaling |
| GO:0014904 | Myotube Cell Development |

**Fig 4.**
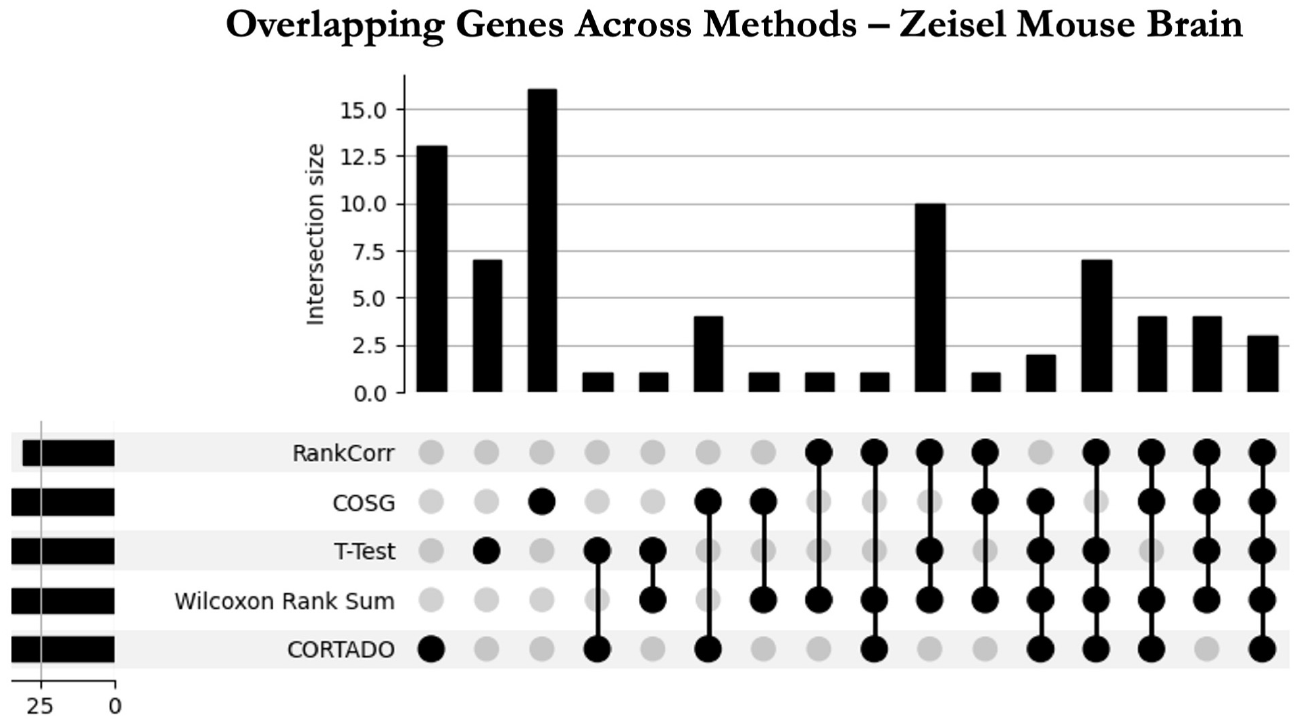
Upset plot depicting overlapping genes selected for each method on Zeisel Mouse Brain Dataset. The horizontal bars on the left represent the total number of genes identified by each method individually, while the vertical bars above indicate the size of the intersections, showing the number of overlapping genes shared among specific combinations of methods. The dots represent the methods included in each intersection, with connected lines linking methods that share genes. The number of overlapping genes for each combination is denoted in the bargraph at the top of the figure.

### CORTADO can identify spatially localized marker genes with biological relevance

We assess the ability of CORTADO to select genes in a spatial dataset context, by analyzing the DLPFC 151507 dataset. The DLPFC 151507 dataset is a spatial transcriptomics dataset derived from the dorsolateral prefrontal cortex (DLPFC), a critical brain region involved in executive functions and working memory. Generated using the 10x Genomics Visium platform, this dataset provides spatially resolved gene expression profiles for approximately 3,639 spots, each linked to specific spatial coordinates within the tissue. It includes information on six cortical layers annotated through histological analysis, enabling the exploration of cellular heterogeneity and tissue organization [25].

The goal of this experiment was to show that the expression of markers selected by CORTADO compared to other methods was more spatially localized, with high expression in the location of interest and low expression in the others. For a more specific comparison, we compared against the two best-performing methods in the benchmarking study we performed, which were the Wilcoxon Rank Sum test and COSG [7] [11]. Similar to the previous analysis, we examined across all clusters the expression of the top 5 marker genes per cluster across methods. In Figure 5A we see the expression of markers in CORTADO and COSG are cell type-specific, with high expression in the cluster of interest and low in all of the others [11]. The Wilcoxon Rank Sum test selects markers with expression in several different clusters, meaning that the markers are not very apt at distinguishing individual clusters. In Figure 5B we see that all methods select markers that have low cosine similarities to other clusters, although to varying degrees of success.

**Fig 5.**
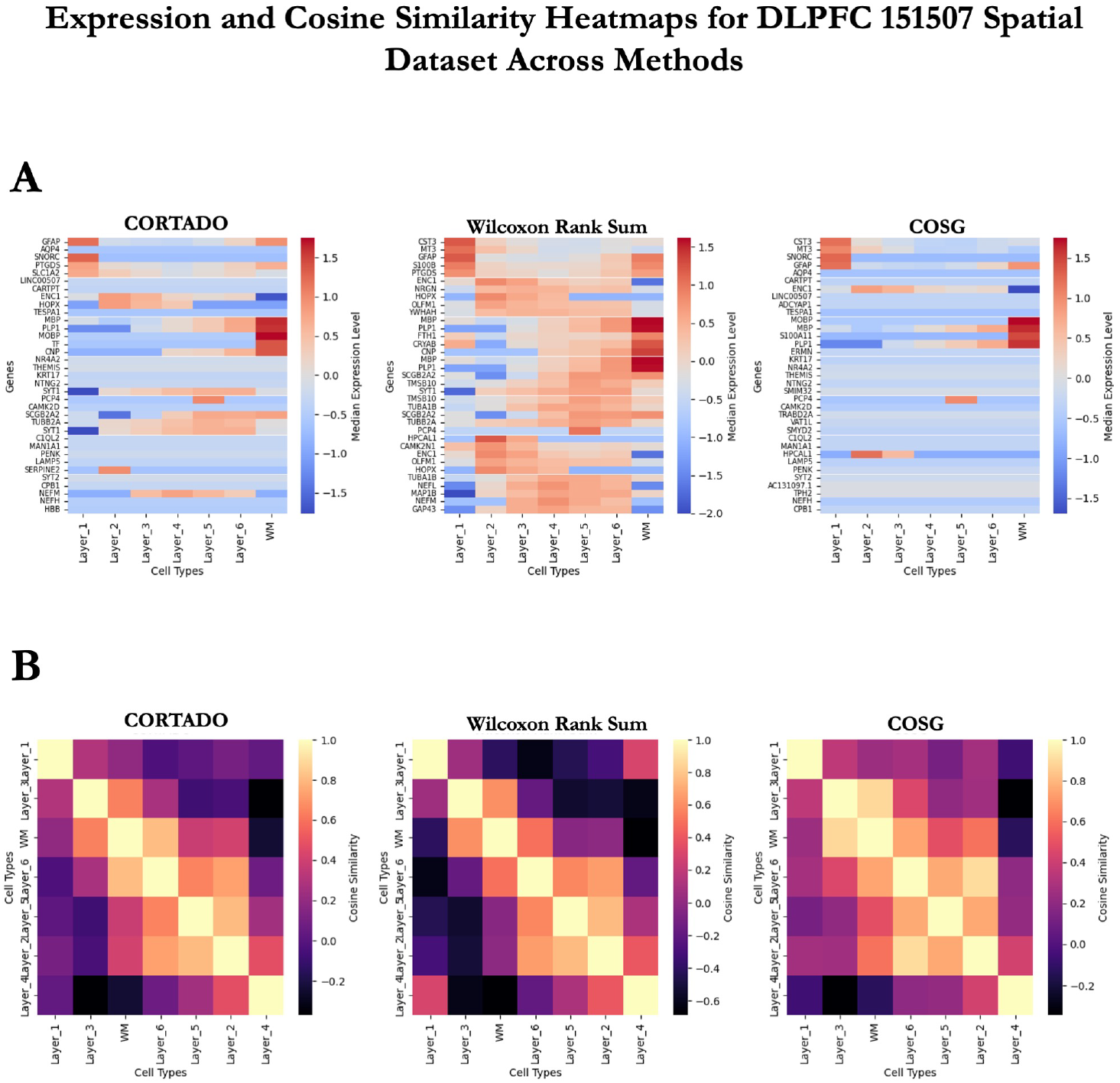
CORTADO analysis on the DLPFC 151507 Dataset. A) Median expression level for top 5 selected markers per method across cell types. B) Cosine similarities between cell-type median gene expression for the top 5 selected markers per method.

When examining the spatial localization-based expression, we looked at the top three markers selected by each method for an example layer, Layer 1. We visualize the localization of expression of these markers in Figure 6. It is clear from this experiment that the markers selected by CORTADO are heavily localized to Layer 1, with the majority of their expression being concentrated there. GFAP and S100B in particular are very specifically isolated to Layer 1. The markers selected by CORTADO compared to the Wilcoxon Rank Sum test and COSG have biological significance. CORTADO selected GFAP as its most significant marker; GFAP is a well-studied marker of prefrontal cortex cells [26]. Abnormal expression of GFAP in the prefrontal cortex can lead to psychotic illnesses such as schizophrenia and bipolar disorder [27]. S100B, which was only selected by CORTADO, is a canonical marker of DLPFC cells categorized by numerous studies. The upregulation of S100B is used to separate DLPFC cells from other cells in the brain, particularly in scRNAseq studies [28] [29] [30]. The Wilcoxon Rank Sum test and COSG on the other hand selected CST3 and MT3, which have high expression across all Layers, and not just Layer 1 [7] [11]. This makes the markers selected by the Wilcoxon Rank Sum test and COSG inherently poor at distinguishing Layer 1 from other clusters. These results demonstrate CORTADO’s superior performance on spatial transcriptomics datasets.

**Fig 6.**
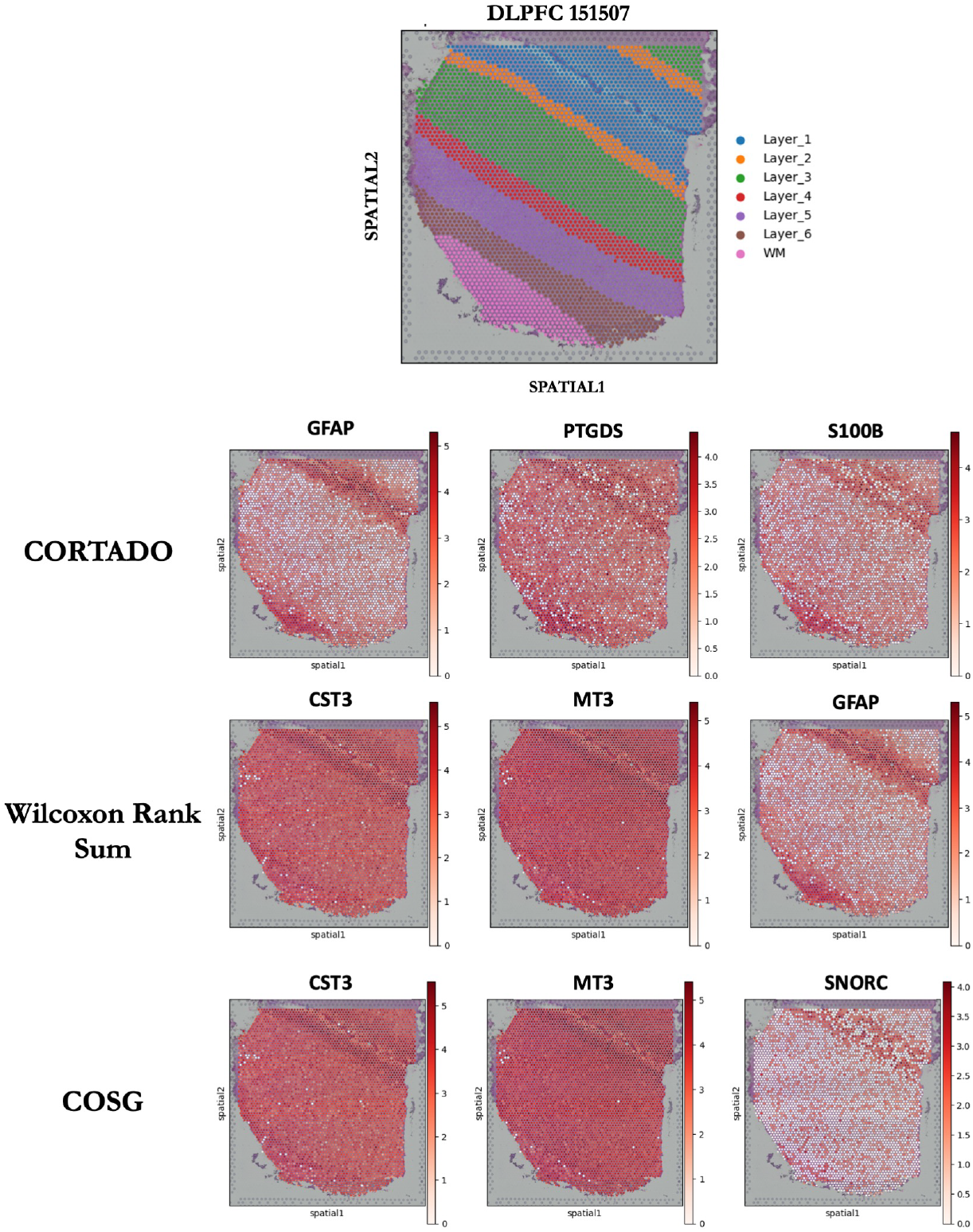
Expression localization of selected markers for CORTADO, Wilcoxon Rank Sum Test, and COSG on DLPFC 151507 spatial dataset.

**Fig 7.**
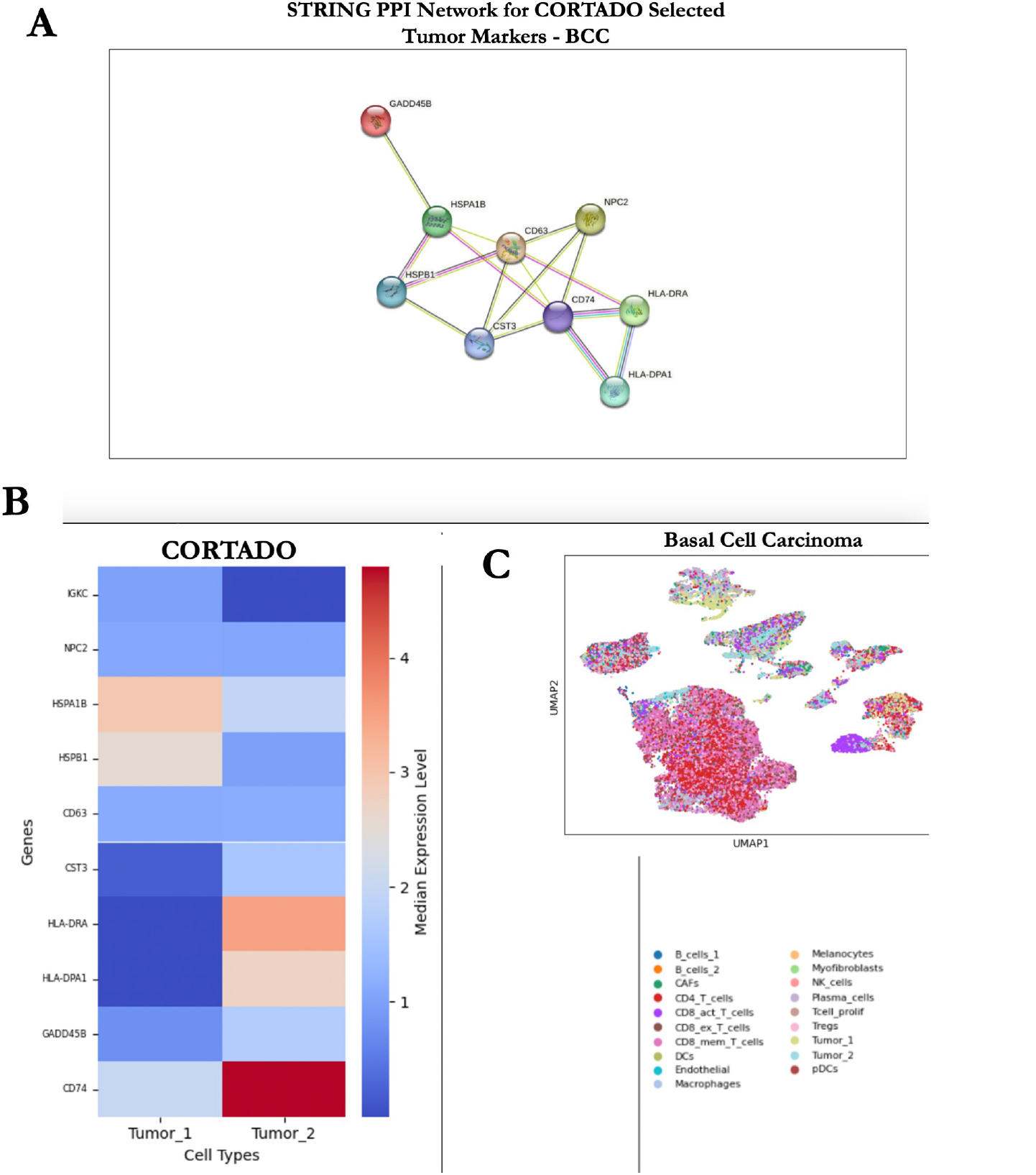
CORTADO Analysis on Basal Cell Carcinoma (BCC) dataset. A) STRING database Protein-Protein Interaction (PPI) network analysis on CORTADO selected markers for Tumor 1 and Tumor 2 clusters. B) Median Expression for CORTADO selected markers in Tumor 1 and Tumor 2 clusters. C) UMAP Visualization for the BCC dataset, which contains 19 clusters. We focused the CORTADO Analysis on the Tumor 1 and Tumor 2 clusters.

### CORTADO reveals novel biomarkers in Basal Cell Carcinoma

The basal cell carcinoma (BCC) dataset contains 53,030 cells from biopsies collected from primary tumor sites of 11 patients with histologically proven advanced or metastatic BCC [19]. This dataset comprises 19 distinct cell clusters, including 2 malignant clusters, and 577 identified genes [19]. To investigate the T cell response to checkpoint blockade immunotherapy (anti-PD-1 therapy) in cancer patients, paired single-cell RNA (scRNA) and T cell receptor (TCR) sequencing were performed [19]. The findings suggest that the efficacy of checkpoint blockade immunotherapy may rely more on newly recruited T cells than on the reactivation of pre-existing T cells within the tumor [19]. CORTADO was applied to the Tumor 1 and Tumor 2 malignant clusters, and the top selected markers for each cluster were analyzed for relevancy to skin cancer.

The top selected markers from the Tumor 1 cluster affect the survival of melanoma cells contingent on their expression levels. HSPA1B, a prognostic biomarker in skin cutaneous melanoma, influences the regulation of the immune response to melanoma cancer cells [31] [32]. Increased expression of HSPA1B correlates with the ability of melanoma cells to evade the immune response, leading to increased melanoma growth and poor overall survival [31] [32]. Another gene within the HSP family, HSPB1, was selected as a top marker that instead inhibits melanoma growth. Cleaving HSPB1 into fragments has a significant effect on tumor progression depending on the fragment [33]. For instance, the secretion of the C-terminal HSPB1 fragment has been found to inhibit the progression of melanoma cells [33]. CD63 also acts as a suppressor of tumor progression, as elevated expression decreases melanoma cell motility and invasiveness [34].

Selected markers from the Tumor 2 cluster play significant roles in the prognosis of melanoma patients. HLA-DRA, a key biomarker linked to the development and progression of skin cutaneous melanoma, ranks as the 46th most strongly associated gene with this disease [35] [36]. HLA-DPA1 was found to be the 80th most closely correlated gene with cutaneous melanoma [36]. Survival analysis showed that increased expression of HLA class II genes, particularly HLA-DP and HLA-DR, in cutaneous melanoma was strongly linked to improved overall survival rates [36]. CD74, a gene that works in conjunction with HLA-DP and HLA-DR genes, is important in the development of adaptive immune responses [36] [37]. It has also been associated with improved prognosis in melanoma patients [36] [37]. This suggests that these selected marker genes may play a critical role in enhancing the immune response against melanoma, contributing to improved patient outcomes. On the other hand, the expression of one selected marker, GADD45B, plays a critical role in the melanoma cell cycle [38] [39]. Down-regulation of this gene can cause accelerated transitions between melanoma cell cycle phases, accelerating tumor growth [38] [39]. CORTADO selected markers that implicate either positive or negative prognosis for melanoma patients, showcasing the model’s use in determining the overall survival of skin cancer patients.

We carry out a protein-protein interaction (PPI) network analysis for tumor biomarkers in basal cell carcinoma to report significant connectivity among nine key proteins (GADD45B, HSPA1B, HSPB1, CD63, CST3, CD74, HLA-DRA, HLA-DPA1, and NPC2), with a total of 16 interactions identified (p-value = 9.14e-07) [40] [41]. The observed number of interactions far exceeded the expected random interactions (*n* = 3), highlighting a significant enrichment of functional associations within this biomarker set. Notably, CD74 emerged as a central hub in the network, interacting with multiple proteins, including HLA-DRA and HLA-DPA1, which are key mediators of antigen presentation and immune response regulation [42]. Other proteins, such as GADD45B and HSPB1, are known to be involved in stress response and cellular homeostasis, which are critical in tumorigenesis and cancer progression [43]. These findings suggest that the identified proteins are not only interrelated but may also contribute collectively to basal cell carcinoma pathophysiology, providing insights into potential therapeutic targets.

The pathways enriched in genes upregulated in tumor-marker cells from the basal cell carcinoma (BCC) cluster primarily involve antigen processing, presentation, and immune activation. These findings provide significant insights into the interactions between tumor cells and immune cells within the BCC microenvironment [19]. The list of pathways enriched is in Table 2. We sorted the pathways in the increasing order of their p-adjusted value meeting the 0.05 cut-off. The upregulation of pathways such as myeloid dendritic cell antigen processing and presentation, antigen processing and presentation via MHC class II, and MHC class II protein complex assembly highlights the role of antigen-presenting cells (APCs), such as dendritic cells, in processing and presenting tumor-associated antigens [44]. These antigens are likely derived from neoantigens generated by UV-induced DNA damage, which is a hallmark of BCC [45]. This activity suggests that tumor cells and associated APCs actively participate in eliciting an anti-tumor immune response by engaging immune cells through MHC class II pathways.

**Table 2.** Enrichment Analysis Results with GO Terms and Pathway Descriptions on CORTADO Markers Selected for both Tumor Clusters in BCC Dataset.

| GO Term | Description |
| --- | --- |
| GO:0002495 | antigen processing and presentation of peptide antigen via MHC class II |
| GO:0019886 | antigen processing and presentation of exogenous peptide antigen via MHC class II |
| GO:0002399 | MHC class II protein complex assembly |
| GO:0002503 | peptide antigen assembly with MHC class II protein complex |
| GO:0002504 | antigen processing and presentation of peptide or polysaccharide antigen via MHC class II |
| GO:0002396 | MHC protein complex assembly |
| GO:0002501 | peptide antigen assembly with MHC protein complex |
| GO:0002478 | antigen processing and presentation of exogenous peptide antigen |
| GO:0019884 | antigen processing and presentation of exogenous antigen |
| GO:0048002 | antigen processing and presentation of peptide antigen |
| GO:0019882 | antigen processing and presentation |
| GO:0050870 | positive regulation of T cell activation |
| GO:0002469 | myeloid dendritic cell antigen processing and presentation |
| GO:0002491 | antigen processing and presentation of endogenous peptide antigen via MHC class II |

**Table 3.** Datasets and their corresponding databases with accession numbers or links.

| Name | Database | Accession Number or Link |
| --- | --- | --- |
| Zeisel Mouse Brain | GEO | GSE103840 |
| Baron Human 1 | GEO | GSE84133 |
| Baron Human 2 | GEO | GSE84134 |
| Baron Mouse 1 | GEO | GSE84135 |
| Baron Mouse 2 | GEO | GSE84136 |
| PBMC3K | 10X Genomics | https://www.10xgenomics.com/datasets/3-k-pbm-cs-from-a-healthy-donor-1-standard-1-1-0 |
| Muraro Pancreas | Broad Institute's Single Cell Portal | SCP2331 |
| BCC Dataset | GEO | GSE123813 |
| DLPFC 151507 | GitHub | https://github.com/LieberInstitute/spatialLIBD |

We also note enrichment in pathways identified by CORTADO, namely, positive regulation of T-cell activation and positive regulation of leukocyte cell-cell adhesion. This indicates that immune cell recruitment and activation are prominent processes in the tumor microenvironment (TME) [46]. This is consistent with the immune-visible nature of BCC, where immune cells infiltrate the tumor to mount a response [47]. The broader pathway enrichment in positive regulation of multicellular organismal processes reflects the systemic impact of localized immune responses within the TME [46]. Signals from tumor cells may propagate to influence systemic immunity or enhance local immune modulation, balancing pro-inflammatory and immune evasion dynamics.

These findings align with the known biology of BCC, a tumor type characterized by its interaction with the immune system due to chronic mutagenic exposure [45]. While immune infiltration and antigen presentation are evident, tumor cells may evade immune clearance through mechanisms such as altered antigen processing, secretion of immunosuppressive cytokines, or checkpoint pathway activation [44].

## Discussion

We introduce CORTADO, a novel method for identifying marker genes in scRNA-seq data. CORTADO combines differential expression analysis with cosine similarity to accurately select discriminative genes. The primary distinction between CORTADO and other methods is its ability to choose genes both based on cosine dissimilarity and differential expression. Moreover, it supports flexible configurations, allowing users to impose constraints on the number of selected markers or to perform unconstrained gene selection, minimizing the number of genes while ensuring their robustness.

In our benchmarking experiment, we found that CORTADO was the highest performer in selecting genes with the highest expression differential, with high gene expression in the cluster of interest versus all other clusters, and was the second-best performer in choosing genes that have lower cosine similarity in other clusters compared to the cluster of interest. CORTADO having a consistently high ranking across both metrics makes it a robust method for selecting strong marker gene candidates.

We also evaluated CORTADO’s performance across multiple datasets. In the case of the Zeisel mouse brain dataset, CORTADO was able to select genes that had strong expression and cosine similarity profiles compared to the other baselines [15]. When examining the markers selected by CORTADO compared to other methods in the case of a specific cluster, Pyramidal CA1 Neurons, CORTADO selected genes that other methods failed to pick up, yet had functional relevance to the CA1 cluster. In the case of the spatial transcriptomics dataset, CORTADO was able to select genes with superior expression localization compared to the baseline methods, COSG and Wilcoxon Rank Sum test [11] [7]. In the PBMC3K dataset, CORTADO was accurately able to identify immune cell markers, which the other methods did not select [18], which we demonstrate in the Supplementary Materials. Finally, CORTADO successfully annotated tumor cell clusters in a skin cancer dataset, with biological relevance validated by existing literature, pathway enrichment, and PPI networks [19].

CORTADO and other commonly used marker selection methods only consider gene expression data to select markers [6]. However, other studies have leveraged multi-omics approaches to determine marker genes, namely DNA methylation data [48]. Future works would include incorporating multi-omics data into the CORTADO framework to capture a more comprehensive marker gene profile. Furthermore, CORTADO inherently selects genes that are highly differentially expressed as an initial filtering step and downselects genes from there. This approach may cause some markers that do not have high differential expression profiles to not be considered due to the initial filtering. Further work would include dimensionality reduction methods to first filter the data by including relevant components, not necessarily purely through expression [49].

## Methods

### Preprocessing

First, a scRNAseq dataset is loaded into Python using the scanpy package. Standard preprocessing steps are completed by filtering lowly expressed genes and performing log-normalization on the raw count data. The log-normalized data is used downstream for the marker gene calculation. The log2foldchange and adjusted *p*-value calculations are performed by the scanpy `sc.tl.rank_genes_groups` function. We use the Wilcoxon Rank Sum test for the calculation of adjusted *p*-value, implemented in Scanpy [7].

### Stochastic-Hill Optimization

#### Constrained and unconstrained hill-climbing optimization

The hill-climbing optimization employed by CORTADO applies a stochastic approach tailored for selecting an optimal subset of genes from a given set. Each element in the binary vector *X* represents a gene, where a value of *X*_*g*_ = 1 indicates that the gene *g* is selected, and 0 means it is not. CORTADO begins with an initial solution and iteratively explores the neighborhood of the current solution to identify better configurations (as per the optimization goals defined next). At each step, neighbors are generated by flipping one or more bits in the binary vector, and their fitness is evaluated based on a predefined objective function. An adaptive exploration mechanism, where the exploration rate decreases exponentially over iterations, balances the need to escape local optima with the focus on refining the solution. The optimization terminates if no improvement is observed for a defined number of steps.

Notably, CORTADO handles both *constrained* and *unconstrained* scenarios for gene selection. In the unconstrained case, *X* is initialized randomly, allowing any combination of genes to be selected without restrictions. In contrast, the constrained case ensures that *X* always contains a fixed number of genes selected (i.e., an exact number of 1s). This constraint is maintained during the generation of neighboring solutions, where any bit-flipping operation must preserve the total count of selected genes. The constrained approach adds complexity but ensures that the optimization process aligns with practical scenarios requiring the selection of a predefined number of genes.

#### Optimization goals

The optimization process for marker gene selection is guided by an objective function with three components, each targeting a specific goal: maximizing the differential marker score within the cluster of interest versus other clusters, minimizing overlap in the expression profiles of selected marker genes, and minimizing the number of selected genes (sparseness). The overall objective function is expressed as:

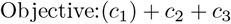

where the components *c*_1_, *c*_2_, and *c*_3_ are defined as follows:

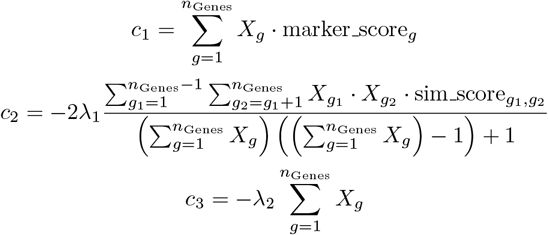

##### 1. Maximizing Differential Marker Scores (*c*_1_)

The term *c*_1_ prioritizes the selection of genes that exhibit a strong differential expression between the cluster of interest and other clusters. For each gene *g*, the marker score is calculated as:

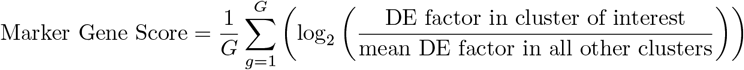

Here, the Differential Expression (DE) factor represents the absolute value of the log2 fold-change for the cluster of interest versus other clusters. By summing the marker scores weighted by the binary variable *X*_*g*_ (indicating whether a gene is selected), this term ensures that the selected genes strongly distinguish the cluster of interest from others.

##### 2. Minimizing Overlap in Expression Profiles (*c*_2_)

The cosine similarity term *c*_2_ penalizes the selection of gene pairs with high similarity in their expression profiles, promoting diversity among the selected marker genes. It is defined as:

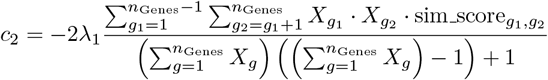

The 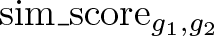 represents the cosine similarity between the expression profiles of genes *g*_1_ and *g*_2_. The term is scaled by *λ*_1_, which determines the relative weight of diversity. By dividing by the number of selected gene pairs, the penalty is normalized, ensuring fair contribution regardless of the total number of selected genes.

##### 3. Encouraging Sparseness in Gene Selection (*c*_3_)

The selection penalty term *c*_3_ discourages the inclusion of too many genes, promoting sparseness in the final selection. It is defined as:

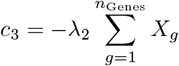

Here, *λ*_2_ controls the penalty strength, with higher values favoring smaller sets of marker genes. This term ensures that the optimization process prioritizes compact and efficient gene sets, which are easier to interpret and validate biologically.

#### Optimization Strategy

The optimization is performed using an iterative procedure that balances these three components. The weights *λ*_1_ and *λ*_2_ are chosen based on cross-validation, ensuring the resulting gene sets meet the goals of specificity, diversity, and sparseness. This approach is particularly effective in single-cell studies, where selecting a minimal set of highly discriminative marker genes is crucial for downstream analysis and experimental validation.

### Benchmarking Methods

#### Scanpy DE Methods: Wilcoxon Rank Sum and t-test

The Scanpy package facilitates differential expression analysis in single-cell datasets, offering versatile methods to identify marker genes across clusters. We implemented differential expression using the `rank_gene_groups` function in Scanpy, modifying the `method` parameter to test with both the Wilcoxon Rank Sum test and the t-test. This flexibility allows for a comparative evaluation of these statistical approaches for detecting differentially expressed genes in single-cell data [7]. The analysis involves calculating the log2foldchange between the cluster of interest and other clusters. Subsequently, the chosen statistical test—Wilcoxon Rank Sum or t-test—is applied to assign a confidence score to each gene, thereby identifying significant markers.

#### COSG

COSG is a standard tool designed for identifying and ranking marker genes, facilitating cell-type classification across various single-cell data modalities. Supporting single-cell RNA sequencing (scRNA-seq), single-cell ATAC sequencing (scATAC-seq), and spatial transcriptomic data, COSG leverages cosine similarity to evaluate gene expression patterns [11]. The method begins by generating an artificial gene that uniquely represents a specific cluster, ensuring it is not expressed in other cell groups. COSG then computes the cosine similarity between the representative vector of this artificial gene and the representative vector of each expressed gene in the dataset. The genes are then ranked based on their COSG scores, allowing researchers to systematically identify high-confidence markers critical for cell-type classification and downstream analyses.

#### scMAGS

scMAGS is a Python-based tool designed for identifying marker genes that define specific cell clusters within single-cell RNA sequencing (scRNA-seq) data. It focuses on selecting genes that best characterize the distinctiveness of cell clusters. The method begins with a cluster-specific gene filtering step, narrowing the pool of potential marker genes by evaluating their relevance to individual clusters [10]. To ensure robustness, scMAGS utilizes quantitative metrics tailored to the dataset’s size and structure. For smaller datasets, the Silhouette index is employed to measure how well a sample aligns with its cluster relative to others. For larger datasets, the Calinski-Harabasz index is used, leveraging its ability to assess the ratio of inter-cluster to intra-cluster similarity. This dual approach allows scMAGS to adapt effectively to varying dataset characteristics.

#### RankCorr

RankCorr is a Python-based tool designed for multi-class marker selection, specifically for single-cell RNA sequencing (scRNA-seq) data. It finds marker genes that distinguish between different cell types or classes [9]. It ranks the input mRNA counts data, converting it into a ranked representation that highlights relative expression levels across genes. Linear separation techniques are then applied to identify distinct patterns within the ranked data, enabling the differentiation of cell types. To ensure adaptability, RankCorr determines the optimal number of markers based on the dataset’s characteristics.

## Data Availability

All datasets used for benchmarking are publicly available. The table below lists the names of each dataset and where they can be accessed.

## Code Availability

CORTADO is available as a PyPi package and hosted on GitHub at https://github.com/lodimk2/cortado-marker.

